# Multiple concurrent feedforward and feedback streams in a cortical hierarchy

**DOI:** 10.1101/2021.01.04.425277

**Authors:** Elham Barzegaran, Gijs Plomp

**Affiliations:** Perceptual Networks Group, Department of Psychology, University of Fribourg, Rue de Faucigny 2, 1700 Fribourg, Switzerland

## Abstract

Visual stimuli evoke fast-evolving activity patterns that are distributed across multiple cortical areas. These areas are hierarchically structured, as indicated by their anatomical projections, but how large-scale feedforward and feedback streams are functionally organized in this system remains an important missing clue to understanding cortical processing. By analyzing visual evoked responses in laminar recordings from six cortical areas in awake mice, we established the simultaneous presence of two feedforward and two feedback networks, each with a distinct laminar functional connectivity profile, frequency spectrum, temporal dynamics and functional hierarchy. We furthermore identified a distinct role for each of these four processing streams, by leveraging stimulus contrast effects and analyzing receptive field convergency along functional interactions. Our results support a dynamic dual counterstream view of hierarchical processing and provide new insight into how separate functional streams can simultaneously and dynamically operate in visual cortex.

## Introduction

Visual processes exhibit complex patterns of fast-evolving activity that are distributed across cortex. Within 100 ms after stimulus onset, activity spreads throughout visual cortex and beyond, both in primates and rodents ^1,2^, enabling functional cortico-cortical interactions that are necessary for even the most elementary visual operations ^3,4^. Such networked processing occurs over dense anatomical projections that reciprocally connect cortical areas. Across visual cortex, the structure of projections indicates a hierarchical organization, with feedforward projections that can propagate sensory activity from lower to higher-level areas, and feedback projections that can exert downstream influences ^5–7^. Understanding how visual cortex enables fast, distributed processing over its fixed hierarchical structure may provide important clues to further understanding large-scale cortical function.

Visual processing critically depends on both feedforward and feedback processes ^8^. Based on onset latencies after stimulation, it was shown that the feedforward spread of activity from V1 mostly follows the structurally defined hierarchy ^1,2^, and that a fast feedforward sweep precedes feedback signals from higher order areas ^9^. But due to the fast mixing of activities after onset, it remains unclear how feedforward and feedback interactions dynamically shape visual processing, which mostly occurs after activity onset.

Candidate mechanisms for how visual cortex could support a mixture of feedforward and feedback processes have been proposed based on the laminar structure of hierarchical projections, and the rhythmicity of neural activity. Recordings across cortical columns have indicated laminar and frequency differences in feedforward and feedback activities ^4,10^, where gamma activity in supragranular layers is thought of as feedforward propagation ^10–13^, and lower-frequency activity in infragranular layers is associated with inter-areal feedback ^14,15^. But although quasi-periodic activity is pervasive in neural systems and much studied, there is no strong consensus on the functional role of different frequency bands ^15–17^. In addition, scale-free activity without a dominant rhythm is an equally widespread phenomenon in neural systems ^18,19^.

To functionally characterize feedforward and feedback streams in the time and frequency domain, we used visually-evoked activity recorded in awake mice across the visual hierarchy with laminar resolution. By using parallel factor analysis of directed cortico-cortical networks, we uncovered the simultaneous presence of two feedforward and two feedback networks, each with distinct laminar connectivity profiles, operational frequencies, temporal dynamics and functional hierarchical organization. We could furthermore determine distinct roles for each of these four processing streams, by leveraging well-described effects of stimulus contrast and by analyzing receptive field (RF) convergency ^20,21^.

## Results

### Evoked activity and functional interactions show complex dynamics

We analyzed local field potentials (LFP) from 11 awake wild-type mice recorded simultaneously from 5 or 6 visual areas with laminar resolution during presentation of high or low contrast drifting gratings (Figure 1a-b). These data were made available by the Allen Brain Institute (see Methods). Stimulus-evoked LFPs showed expected fast dynamic responses across layers and areas throughout the cortical hierarchy (Figure 1c) ^2^, in line with results from non-human primates ^1^.

**Figure 1.**
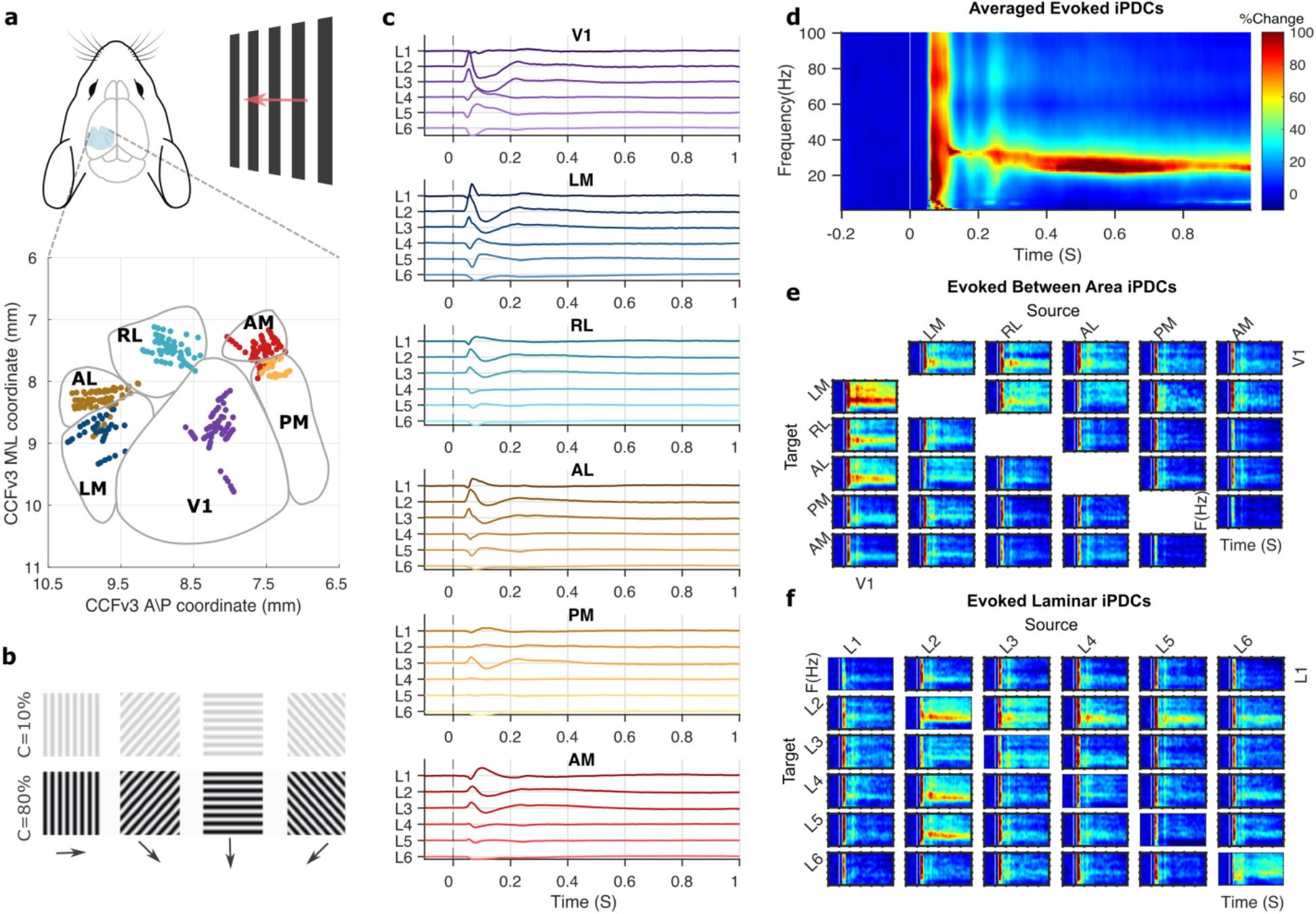
Visual stimuli evoke large-scale activity and functional interactions. **a,** Laminar recording sites across cortical areas from 11 mice, in Allen brain institute’s standard CCFv3 space. **b,** Visual stimuli were drifting gratings of high (80%) or low (10%) contrast, presented for 2 S. **c,** Grand-average bipolar LFPs evoked by high contrast stimuli, for L1 to L6 in six visual areas, shown in anatomical hierarchical order ^7^. **d,** Directed functional connectivity (iPDC) in the time and frequency domain, averaged over all areas, layers, and animals. **e,** Time-frequency connectivity between areas, averaged over layers and animals. **f,** Time-frequency connectivity between layers, averaged over areas and animals. Heatmaps show percentage of post-stimulus change, columns indicate sources and rows indicate targets of directed functional interactions.

To derive functional interaction strengths between all areas and layers, we used an optimized Kalman filter to model dependencies between LFPs and calculated the information Partial Directed Coherence (iPDC) ^22,23^. iPDC provides a multivariate measure of directed functional connectivity (Granger causality) with high temporal and frequency resolution. The resulting connectivity matrices revealed a detailed time-frequency representation of between-area interaction strengths across visual cortex, with laminar resolution (Figure 1d-f).

Averaging across areas and layers, we found that stimulus onset evoked statistically robust functional connectivity, and that the interactions were contrast-sensitive. Evoked connectivity onset was around 50 ms for high contrast, and 85 ms for low contrast stimuli (Figure 2), following known contrast dependency of LFPs ^20^. The initial response was followed by sustained, frequency-specific increases in gamma band interactions that peaked around 35 and 25 Hz for high and low contrast stimuli respectively. This indicates that reducing stimulus contrast lowers the gamma band interaction frequency, similar to its effect on LFP power spectra ^16,24,25^.

**Figure 2.**
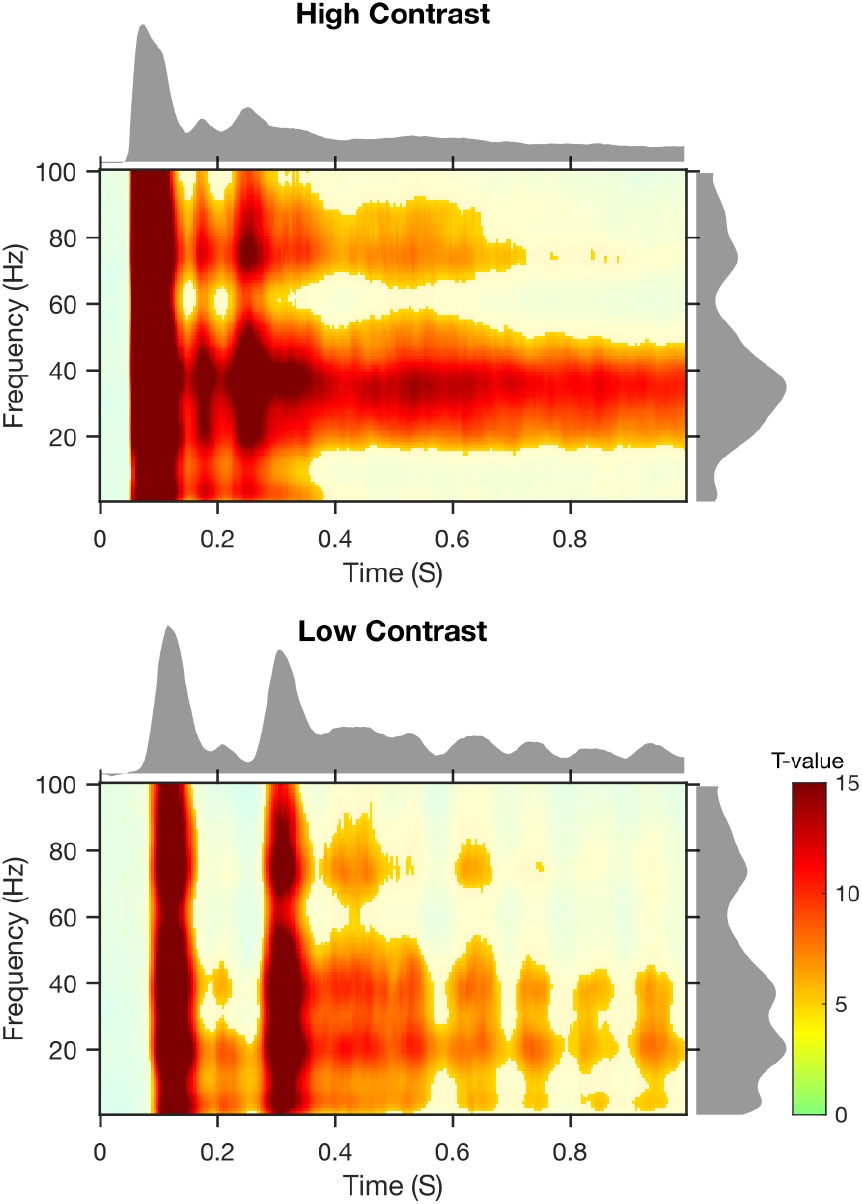
Visually-evoked functional connectivity for high and low contrast stimuli. Time-frequency plots show statistically significant functional connectivity increases with respect to baseline (one-way t-tests *p*<0.05, Bonferroni corrected), marginal distributions reflect t-values. Data averaged across layers and areas.

### Four concurrent networks in mouse visual cortex

To further analyze the time- and frequency-resolved connectivity across visual cortex, we used Parallel Factor Analysis (PARAFAC), a tensor rank decomposition method that provides robust and interpretable results ^26^. PARAFAC has previously been used to identify constituent components of time-varying power spectral density (PSD) ^27,28^ and functional connectivity ^29^. PARAFAC guarantees a unique solution when the appropriate number of components is selected. To select this number, we combined multiple criteria, including the core consistency diagnostic, mean square error, model convergence, and visual inspection ^26,28,30^. This showed that the functional connectivity matrices were best decomposed into four components, independently for low and high contrast conditions, and explaining around 90% of total variance. We confirmed the reliability and validity of the four-way model across animals using a bootstrap analysis that showed high consistency across bootstraps for each component (average correlation coefficients for all components > 0.81, *p*<0.05), and low correlations between components (average correlation coefficients = 0.21, not significant).

This way, we established that the dynamic laminar functional connectivity pattern across the visual hierarchy can be decomposed into four distinct networks, each of which reflects a processing stream defined by its directed connectivity strengths between areas over source layers, target layers, frequency spectrum and temporal dynamics (Figure 3).

**Figure 3.**
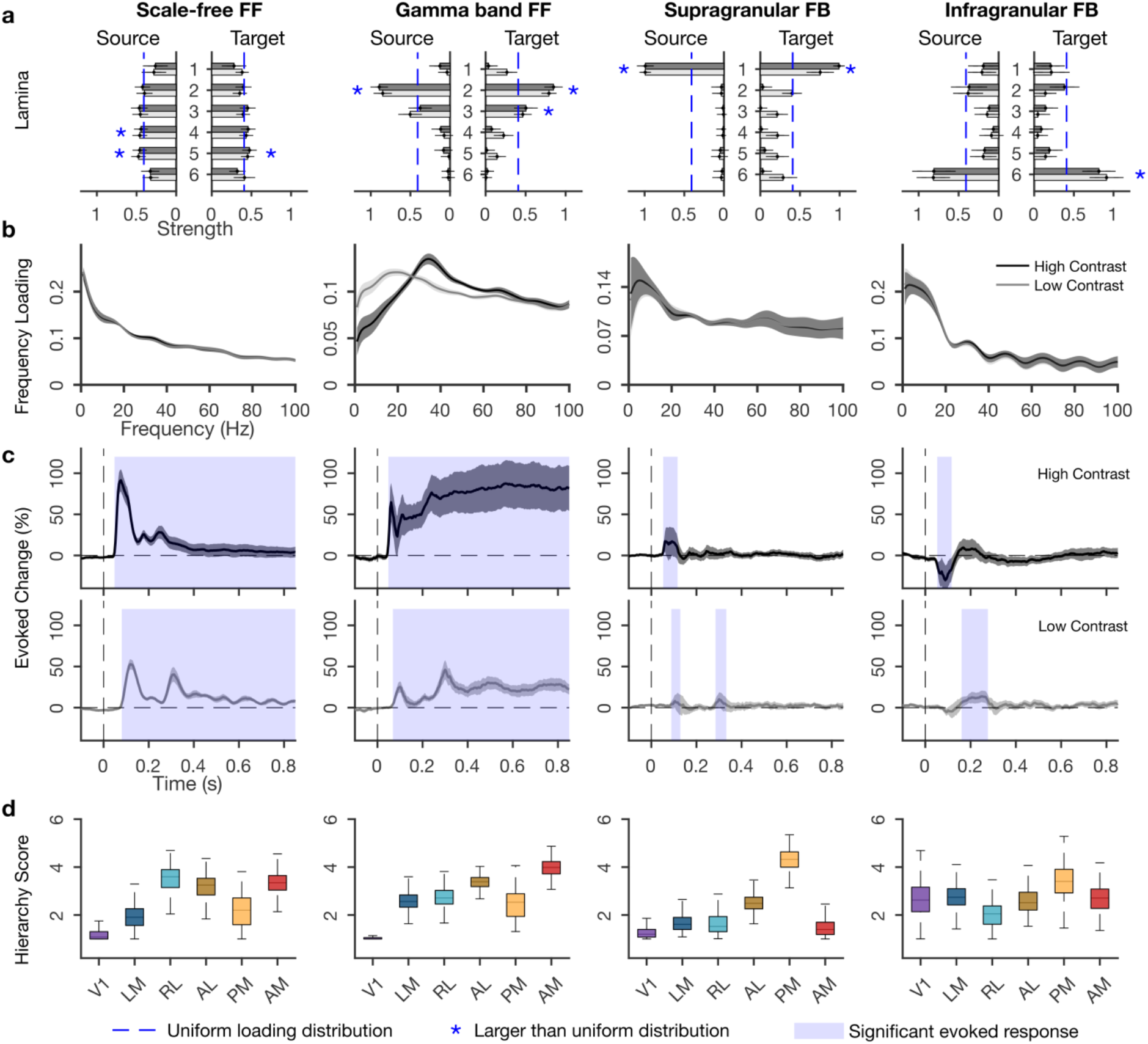
Four concurrent functional networks in visual cortex. **a,** Laminar input and output strengths (loadings) for the two feedforward (FF) and two feedback (FB) networks. Blue stars indicate layers with larger amplitude than the uniform distribution, calculated based on 95% confidence interval. **b,** Frequency distributions of each network. **c,** Temporal dynamics per network, as percentage of change compared to pre-stimulus period, calculated from temporal loadings. Shading indicates statistically significant increases from baseline. **d,** Functional hierarchy scores calculated from between-area connectivity loadings (see Methods). The higher and lower limits of box plots indicate bootstrapped 25 and 75 quantiles of the hierarchy scores, whiskers indicate extreme values excluding outliers. For a, b, and c error bars and grey shading indicate standard deviations over bootstraps (n=500).

### A dominant scale-free network

We found that a scale-free network was the dominant network, accounting for about 50 ± 5 % (mean ± s.d.) of the PARAFAC model amplitude, in both stimulus contrasts. The connectivity strengths in this network were relatively uniformly distributed over source and target layers, but showed significantly stronger connections from L4 and L5, and toward L5 than expected by chance (Figure 3a). This pattern of laminar connectivity resembles the distribution of feedforward targeting neurons through infragranular layers, particularly L5, as established by retrograde tracing in mouse ^7^ and monkey ^6^.

The network’s frequency spectrum followed a power-law distribution for both low and high contrast stimuli, revealing a scale-free temporal dynamic where no particular rhythm dominates the functional interactions (Figure 3b). Power-law distributions take the form 1/f^β^, where *β* quantifies the self-similarity in the time domain (autocorrelation). Model fitting of the scale-free network showed a *β* exponent of 0.4, lower than typical autocorrelations for electrophysiological data, which fall in the range of 1-3 for the frequency range of 1-100 Hz ^19^. To test whether the low *β* values are characteristic of the network or of the underlying LFPs, we fitted power law distributions to the PSDs of LFPs in all areas. This revealed higher *β* exponents (2 ±0.5 mean ± s.d. over bootstraps), demonstrating that low self-similarity rather is a property of the scale-free network than of the underlying LFPs. These results indicate that between-area functional interactions have greater agility in time than the within-area activity.

The scale-free network showed robust amplitude increases quickly after stimulus onset, which decreased after around 150 ms, but exceeded pre-stimulus values throughout the epoch (*p*<0.05, Bonferroni corrected, Figure 3c). This fast network activation, together with its laminar profile supports the idea that sensory activation is quickly relayed across areas over infragranular layers ^31^. The scale-free network showed different dynamics depending on stimulus contrast. With low contrast, onset latencies were delayed (from around 50 to 80 ms) and a more pronounced second peak occurred at around 300 ms before amplitudes decreased (c.f. Figure 2).

To characterize whether the direction of interactions followed a predominant feedforward or feedback pattern, we derived functional hierarchy scores from between-area connectivity strengths ^12^ (see Methods). This revealed a hierarchical organization with V1 and LM at the bottom followed by the other areas (Figure 3d), indicating a mainly feedforward direction of interactions. The functional hierarchy resembled the structural hierarchy established from axonal projections ^2,7^, but it put PM at a lower ordinal position. Area PM, like V1, preferentially responds to low temporal and high spatial frequencies, while LM, AL, RL, and AM prefer high temporal and low spatial frequencies ^32^. The temporal and spatial frequencies of the drifting gratings were closer to the preferred frequencies of PM, allowing it to more strongly drive activity in other areas, thus lowering its position in the functional hierarchy.

In summary, we found a dominant scale-free network whose laminar connectivity profile, and functional hierarchy indicate a predominant feedforward direction of interactions. The network responds strongly but transiently to stimulus onsets with some contrast sensitivity.

### Visual stimuli recruit a feedforward gamma band network

The second strongest network contributed 18 ± 2% of total model amplitude, for both stimulus contrast. Its laminar connectivity pattern showed L2 and L3 as both the strongest source and target layers (Figure 3a), a pattern that is typically associated with feedforward processing streams in mouse and primate ^6,33–35^.

This network showed clear peak amplitudes in the gamma band, broadly defined as 25-100 Hz. Visually induced gamma band activity is typically linked to feedforward processes ^11,24^ and may help synchronize activities between cortical regions ^13^. The network’s peak frequency changed with stimulus contrast, centering around 38 Hz for high contrast and around 26 Hz for low contrast (Figure 3b), resembling the frequency shift observed in overall functional connectivity strengths (Figure 2), and the contrast sensitivity of gamma band LFP responses ^16,25^. Such strong contrast-sensitivity indicates a role in processing stimulus content.

The gamma band network also showed contrast-dependency in its temporal dynamics (Figure 3c). For high contrast stimuli, this network showed increased amplitudes after 50 ms that were sustained throughout stimulus presentation. In the low contrast condition, amplitude increase started around 70 ms, and were sustained with an apparent rhythmic modulation. These effects resemble the contrast effects in overall functional connectivity and those in the scale-free network.

Functional hierarchy analysis confirmed the feedforward character, with V1 located lowest and the ordinal position of the other areas following the structural hierarchy ^7,34^. As in the scale-free network, the gamma band network showed a relatively low hierarchical position for area PM, possibly due to the stimulus characteristics and response properties of this area ^32^.

In sum, the gamma band network shows a laminar inter-areal connectivity pattern and hierarchical organization that conforms to known feedforward structural connectivity patterns. The network showed a sustained response to visual stimulation that strongly depended on stimulus contrast.

### Two low-frequency feedback networks

In addition to the feedforward networks, we uncovered two feedback networks that accounted for 17 ± 2% and 15 ± 4% of model amplitude. One was a supragranular network, with putative L1 as the main source and target of inter-areal connections, the other was an infragranular network with L6 as the main source and target (Figure 3a). Anatomical studies of primate visual cortex in monkey established that besides the well-known feedback pathway over L6, another feedback pathway exists over supragranular layers L1 and L2 ^6,34,35^. Our findings for the first time distinguish these two feedback pathways using analysis of functional data and demonstrate their presence in mouse.

In accord with a presumed feedback function, we found that both networks showed a broad low-frequency distribution, which peaked at 5 and 6 Hz respectively and included the alpha band (full-width half-maximum of 1-14 Hz and 2.5-15.5 Hz respectively). Previous works identified low-frequency interactions as signatures of feedback processing in visual cortex of cat and monkey, using functional connectivity analysis and causal interference ^10,36^.

The two feedback networks showed strikingly different evoked responses and contrast sensitivities. The supragranular feedback network showed fast and transient evoked amplitude increases with typical contrast sensitivity, i.e. slower onset and reduced amplitudes for low contrast stimuli (50 vs. 85 ms). In addition, low contrast stimuli evoked a second amplitude increase at around the same latency that the feedforward networks showed a second low-contrast peak. The infragranular feedback network, conversely, showed quickly decreased amplitudes for high contrast stimuli, between 50 and 120 ms, but increased amplitudes for low contrast stimuli at longer latencies, between 155 and 280 ms. This indicates enhanced feedback for low contrast stimuli at longer latencies ^9,21^. In addition, these findings show how stimulus contrast differentially recruits the two feedback pathways, first over infragranular and then over supragranular pathways, suggesting a functional dissociation of feedback streams.

The feedback networks had distinct functional hierarchies. The supragranular network showed AM and V1 at the bottom of a hierarchical organization that otherwise resembled the structural hierarchy ^2,7^, but with lower hierarchy scores for RL and AL than in the feedforward networks. The low hierarchical positions of high-level visual areas characterize the supragranular network as a feedback network, in line with its laminar connectivity and frequency spectrum. For the infragranular network, area RL showed the lowest hierarchy score, but the other areas appeared not to be hierarchically ordered, with V1 having a higher hierarchy score than in any of the other networks. The low RL position supports a predominant feedback directionality for the infragranular network.

Taken together, these findings distinguish two feedback networks, one operating over supragranular and one over infragranular layers. The networks share a low-frequency profile but differ in how they respond to stimulus onset, their contrast sensitivity, and their functional hierarchies. This differentiation is in line with the distinct influences of feedback arriving in the superficial or deep layers, and how this affects neural activity in the cortical column ^31,37–39^.

### Rhythmic modulation of feedforward networks

The feedforward networks appeared to show rhythmic amplitude modulations in time, especially for low contrast stimuli (Figure 3). To investigate this, we applied Fourier transformation on the post-stimulus window of temporal loadings, after removing linear trends. This revealed a rhythmic modulation of both feedforward networks that was most prominent for low contrast stimuli and had a peak frequency of around 5 Hz (Figure 4).

**Figure 4.**
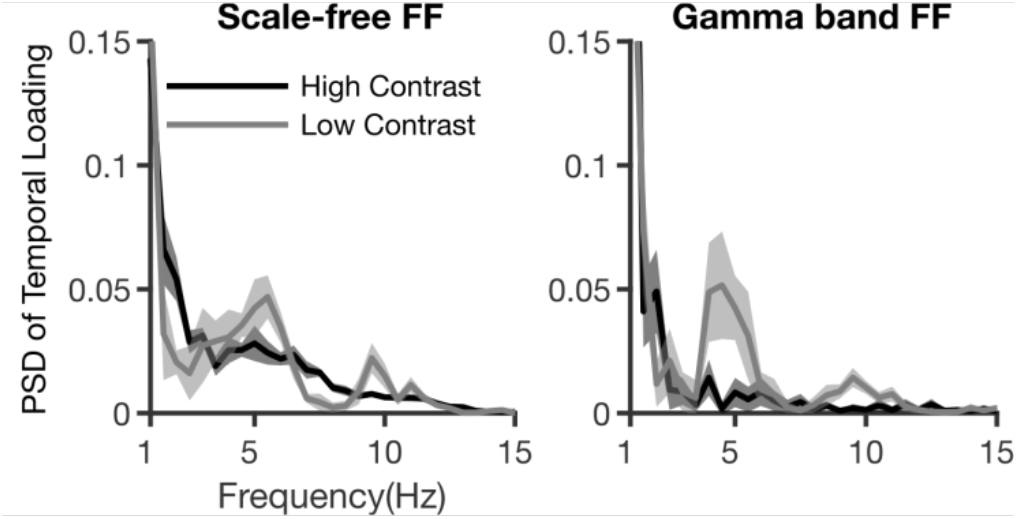
Spectral power of feedforward network dynamics. Fourier transformation of temporal dynamics for feedforward networks, for two contrast conditions. Shaded area indicates standard deviation over bootstraps.

Theta-band rhythms in mice are common in hippocampus, but also occur in sensory areas ^15,40^. In the current data, the theta modulation is likely of cortical origin because it varies with stimulus contrast. The possibility that micro-saccades account for the theta modulation of feedforward networks was excluded in a control analysis (data not shown). Theta rhythms in sensory cortex have been shown to organize local activity, in mouse and monkey ^40–42^. Although theta-rhythmic functional interactions were previously reported in monkey ^43^ and cat ^14^, our results show a rhythmic amplification of entire feedforward networks, particularly when stimuli are less visible.

### Receptive field distances and functional connectivity

Considerable evidence shows that feedforward projections tend to converge onto matching receptive fields (RF) in upstream areas, whereas feedback projections tend to be more divergent 6,8,44. Convergent and divergent projections play different roles in spatial processing, and we therefore asked how inter-areal functional connectivity strengths related to the distance between RF centroids in the source and target location. We hypothesized that functional connectivity strengths would increase with RF overlap for feedforward, but not for feedback networks.

To test this, we modeled functional connectivity strength as a function of distance between RF centroids for all recording locations (areas and layers, see Methods). This confirmed the convergent nature of interactions for the gamma band feedforward network (*p* < .001, 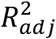 = .88), but not for the scale free network (*p* = .26, 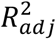 = .76) (Figure 5). In addition, this analysis uncovered a divergent pattern of interactions for the infragranular feedback network, with stronger interactions toward more spatially distinct RFs (*p* < .001, 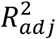 = .89). No RF-dependence was observed for the supragranular feedback network (*p* = .16, 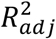 = .7). These results reveal spatial processing differences between the feedforward and feedback networks by showing how the spatial organization of RFs in the source and target location co-determine functional interaction strengths.

**Figure 5.**
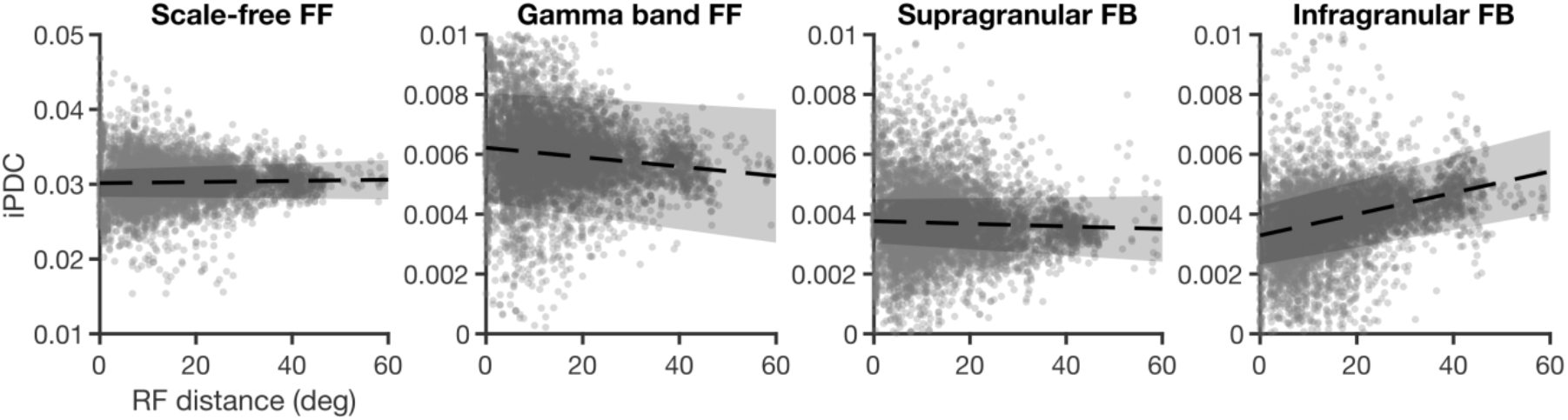
Connectivity as a function of RF distance. Dots represent values acrossss bootstraps, dashed line represents the regressed line and the shaded area indicate the 95% confidence intervals (on intercept and slope of the model). Data from high contrast condition.

## Discussion

We found that functional interactions across visual cortex can be reliably decomposed into four constituent directed networks, based on their distinct laminar connectivity profiles, operational frequencies, temporal dynamics, and hierarchical organizations. The laminar connectivity pathways match anatomical knowledge and provide strong functional evidence for the simultaneous presence of multiple feedforward and feedback streams ^6,33^, and extend the notion of functional hierarchies in cortex ^2^. We furthermore show distinct functional roles for each of these streams in response to stimuli, reconciling the hierarchical structure of visual cortex with its fast distributed processing.

Our analyses demonstrate the existence of two distinct feedforward processing streams. Previous studies have associated feedforward sensory processing with gamma band activity in supragranular layers ^10,12^, in line with anatomical connectivity patterns of these layers ^6,34^. Our gamma band network provides strong functional support for a feedforward role for layers 2/3 pathways mediated by gamma band interactions. But in addition, our results indicate a second feedforward stream over infragranular layers that has a scale-free frequency distribution. Evidence of an infragranular feedforward pathway has been previously reported based on anatomical data ^6^, but to our best knowledge it has not been reported in functional studies.

Our results identify distinct roles in visual processing for the scale-free and the gamma band network. An important clue to the possible function of the scale-free network lies in its power-law frequency distribution. Previous works have related scale-free LFP spectra to overall neuronal population activities ^45^, but at a larger-scale, power-law distributions can indicate that a complex system is working near criticality, i.e. at an equilibrium point between two states ^18^. Dynamics near a critical point can allow neural activity states to rapidly adjust to changes in the external environment, by providing a flexible mid-level synchronization between neuronal rhythms and fast transition between states via phase resetting ^46^. In our data, the scale-free network was the dominant network (strongest amplitude), suggesting its relative importance and a possible role in facilitating the state and activity changes of other frequency-specific networks in response to visual stimulation, an interpretation that is supported by its strong but transient response to visual stimuli, and its lack of RF convergence. In line with such a dynamic resetting role, the scale-free network also showed low autocorrelation in time, and non-convergent RF projections of its upstream interactions, indicating nonspatial processing.

The gamma band network, by contrast, responded to visual stimuli with a sustained amplitude increase throughout stimulus presentation. This suggests a role in processing stimulus content that is supported by its sensitivity to stimulus contrast, both in the time and frequency domain. In addition, the RF convergence of its interactions make it suited to process spatial stimulus aspects, because convergent interactions preserve spatial position while propagating induced activity up the hierarchy. The peak frequency and amplitude of the induced gamma oscillations are previously reported to be modulated by stimulus properties such as contrast, size and location 11,16, as well as by selective attention ^47^. Overall, it has been shown that salient or attended stimuli induce faster and stronger gamma oscillations in early visual areas, and when competing gamma oscillations exist within low-level areas, the faster oscillations have a higher chance to induce activity in higher-level areas through phase coding ^13,25^. Therefore, the stimulus sensitivity of gamma oscillations has been described as an implementation of bottom-up attentional selection ^13^. In our results, the contrast sensitivity of gamma peak and amplitude indicate that elementary forms of such an effect might exist in mice.

We found strong evidence for a theta-band modulation of the feedforward networks. This theta modulation seems unlikely to be caused by the feedback networks, because they did not show a sustained response, whereas the theta modulation persisted over the whole period of stimulus presentation (Figure 4). Given its contrast dependency, a cortical origin seems the most likely. Cortical theta activity in sensory areas has been linked to feedforward interactions ^12,17^, and plays a role in rhythmic perceptual sampling in monkey and human ^41,48^. A proposed mechanism of perceptual sampling is based on theta-gamma cross-frequency coupling, where theta modulates feedforward activity through resynchronization of gamma oscillations between lower and higher visual areas by resetting of the gamma phase ^13,42^. Our results show that entire feedforward networks can be modulated by a theta rhythm and provide first evidence for a rhythmic visual sampling in mouse. If behaviorally verified, this means that rhythmic visual sampling is an evolutionary well-preserved feature, and that the causal investigations possible in mice can be employed to help further understand this phenomenon.

We uncovered two feedback networks that operate at alpha and theta frequencies ^10,12^. Yet each feedback network showed distinct laminar connectivity, functional hierarchy, contrast sensitivity and relationship with RF distances (Figure 3), suggesting a functional dissociation between feedback streams for stimulus-dependent spatially-organized feedback and global modulatory feedback ^8^. The infragranular feedback stream showed a particular contrast sensitivity, with fast amplitude decreases for high contrast, and slower but amplitude increases for low contrast stimuli. In addition, its connectivity pattern was spatially organized, with stronger connections to locations with nonoverlapping RFs. This organization is compatible with a role in contextual spatial processing, such as surround suppression and figure-ground segmentation ^4,10,44^, operating over infragranular layers in stimulus-specific ways ^49^. It has been shown that feedback over infragranular layers can affect local processes in the cortical column through upward propagation ^31,37^. Likewise, feedback at supragranular layers can modulate activity throughout the column ^37^. In our data, the supragranular feedback stream showed no spatial organization of connections, and less sensitivity to contrast, making it suitable for global or non-retinotopic feedback modulations ^50^. In line with this characterization, the lowest areas in its hierarchy were motion-sensitive areas AM, LM and RL ^32^.

Feedback processing is generally thought to arise after a first feedforward sweep, in a two-stage process ^9^. Our results show partial support for such a sequential view, in that infragranular feedback increased at longer latencies (150 ms), and selectively for low contrast stimuli where feedback is expected to play a greater role ^21^. In addition, however, our results show that stimuli can evoke quick feedback processing over the supragranular stream, and that they can also quickly suppress ongoing feedback processes. Suppression of theta and alpha feedback due to attention has been previously reported ^42^. Taken together, our results characterize visual-evoked responses as a dynamic re-weighting of feedforward and feedback processes, which are better separable in terms of their functional hierarchies, laminar connectivity and operational frequencies, than they are in time.

Overall, feedback amplitudes were small compared to the feedforward amplitudes ^35^, and while feedback was transient, feedforward networks showed more robust evoked responses and for longer time. Anatomical studies in monkey show that feedback projections are more numerous than feedforward projections, but that feedforward projections are stronger, especially at shorter range ^6^. Our results confirm this with functional evidence and characterize the evoked response as a predominantly feedforward process. But even though the stimuli and task were not optimized for investigating typical roles attributed to feedback connections, like figure-ground segregation ^4,21^ or contour integration ^3^, the demonstrated presence of two feedback streams in awake but passive mice points to a fundamental importance for feedback in vision. It also reveals similarities between the large-scale functional organization of processing streams in mouse and primates which open up new avenues for investigating large-scale cortical function using causal manipulations.

## Methods

### Dataset and visual stimuli

We used publicly available recordings from the Visual Coding - Neuropixels dataset, provided by the Allen Institute for Brain Science ^2^. This dataset contains LFPs recorded simultaneously in 4 to 6 visual areas of awake mice, using Neuropixel probes (^51^, 40 μ*m* distance between recording channels), with 2.5 KHz sampling rate. Full details on surgery, stimulation protocols and recording techniques are available in the technical white paper ‘Allen Brain Observatory – Neuropixels Visual Coding’, 2019 (portal.brain-map.org/explore/circuits/visual-coding-neuropixels). The data was downloaded in the format of raw Neurodata Without Borders (NWB), using Allen SDK 1.2.0 (2019).

We used a subset of the Brain Observatory stimulus set with a large number of trials, the Functional Connectivity stimulus set. From 14 available LFP recordings in wild type mice, we selected 11 for analysis (average age = 126 ± 10 days, 10 males) that showed clear source and sink patterns of current source density (CSD) and had simultaneous recordings from at least four separate areas (Figure 1a). These areas were V1 (recordings from 11 animals), AL (10), RL (9), PM (7), LM (6), and AM (11) of the left hemisphere; 2 animals had simultaneous recordings in 6 areas, 6 in 5 and 3 in 4.

From the standardized battery of visual stimuli ^52^, we selected drifting gratings for their reliably evoked signals and high number of trials (75 trials for each grating orientation and contrast). Gratings were presented in random order, with a high (80%) or low (10%) contrast at detectable levels ^53^, four different orientations (0, 45, 90 or 135 degrees), and with a spatial frequency of 0.04 cycles/deg and temporal frequency of 2Hz (drifting left-to-right). Stimuli were presented monocularly to the right eye on a monitor that covered 120 × 95 degrees of visual angle. The paradigm consisted of two seconds of stimulus presentation, and one second of inter-stimulus interval during which a uniform mean luminance gray was presented.

### Preprocessing and layer assignment

For each trial, we considered the epoch from 300 ms before to 1000 ms after stimulus onset. We pooled the epochs from four orientations together which resulted in 300 epochs per animal, per contrast.

To identify representative electrodes for each cortical layer (L1 to L6) we derived CSD patterns from averaged LFP data ^54^. We assigned layers based on sources and sinks across cortical depth, following ^40^, taking into account depth estimates provided with the dataset. The average cortical depths of selected electrodes for each layer are listed in Table 1.

**Table 1.**
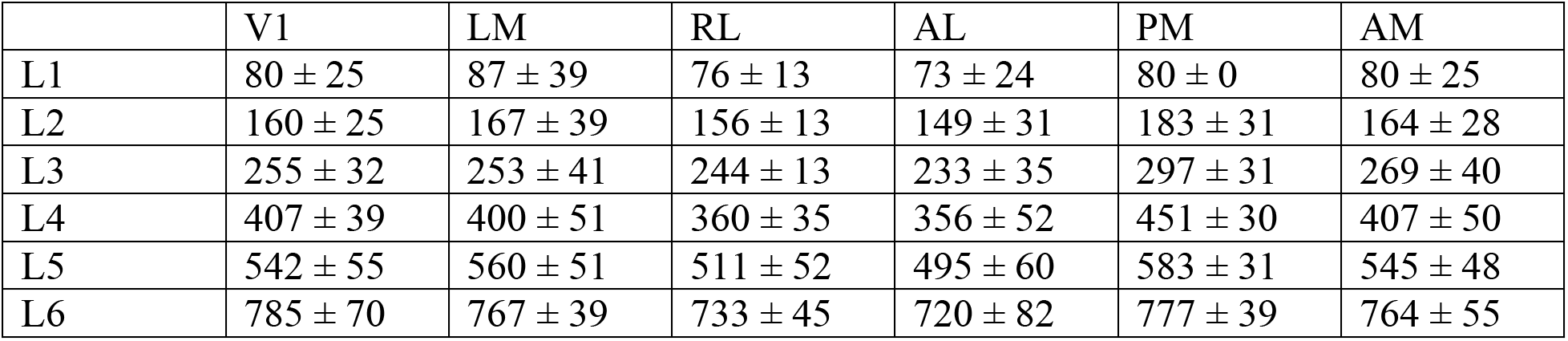
Average cortical depth of selected channels for each layer and area (mean ± standard deviation)

|  | V1 | LM | RL | AL | PM | AM |
| --- | --- | --- | --- | --- | --- | --- |
| L1 | 80 ± 25 | 87 ± 39 | 76 ± 13 | 73 ± 24 | 80 ± 0 | 80 ± 25 |
| L2 | 160 ± 25 | 167 ± 39 | 156 ± 13 | 149 ± 31 | 183 ± 31 | 164 ± 28 |
| L3 | 255 ± 32 | 253 ± 41 | 244 ± 13 | 233 ± 35 | 297 ± 31 | 269 ± 40 |
| L4 | 407 ± 39 | 400 ± 51 | 360 ± 35 | 356 ± 52 | 451 ± 30 | 407 ± 50 |
| L5 | 542 ± 55 | 560 ± 51 | 511 ± 52 | 495 ± 60 | 583 ± 31 | 545 ± 48 |
| L6 | 785 ± 70 | 767 ± 39 | 733 ± 45 | 720 ± 82 | 777 ± 39 | 764 ± 55 |

In order to remove the common reference, which could cause spurious estimation of functional connectivity, we applied bipolar re-referencing by computing the difference of the LFPs from two neighboring channels ^55,56^. LFP data were downsampled to 250 Hz after anti-aliasing filtering.

### Functional connectivity

To estimate time- and frequency-resolved directed functional connectivity we used a multivariate parametric approach, the Self-Tuning Optimized Kalman filter (STOK) ^23^. STOK uses a Kalman filter formulation that is optimized to model rapidly fluctuating between-signal dependencies under unknown noise conditions, resulting in a time-varying multivariate autoregressive (tvMVAR) model. To estimate the tvMVAR model, a model order parameter *p* should be estimated prior to analysis. This parameter indicates how much of the past information should be included in estimation of current state. Here we selected the optimal model order by minimizing the difference between the tvMVAR model power spectra and the spectra calculated by a nonparametric method based on complex Morlet Wavelet ^57^, for each animal separately. We then selected the maximum optimal order observed over animals (*p* = 15 samples, or 60 ms) as the model order, and calculated tvMVAR models per animal and condition (low, high contrast) using all single trial epochs and a filter factor of 0.98. STOK code is freely available for Matlab (https://github.com/PscDavid/dynet_toolbox) and Python (https://github.com/joanrue/pydynet).

After Fourier transformation, tvMVAR coefficients were normalized to express a directed measure of functional connectivity in line with Granger causality, the information partial directed coherence (iPDC) ^22,59,60^. Resulting iPDC matrices contained the layer-specific directed functional connectivity between all recorded areas, for every frequency between 1 and 100 Hz (1 Hz resolution), and timepoint between −300 to 1000 ms around stimulus onset (4 ms resolution). We preserved the between-area connections for further analysis, averaged across animals, and unfolded the resulting connectivity matrix into 5 dimensions as follows: source layers, target layers, time, frequency, and between-area connections (30 directed connections between the 6 areas).

### Parallel Factor Analysis

PARAFAC decomposes a multiway matrix into a fixed number of components, with each component represented by a set of loading vectors that correspond to the original data dimensions ^26,61^. We applied PARAFAC to the averaged 5-dimensional connectivity matrices. The resulting PARAFAC decomposition can be expressed as:

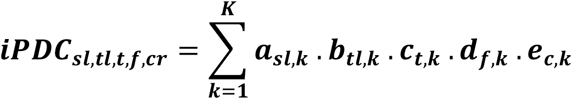

Where K is the number of components, and *a*_*sl*,*k*_, *b*_*tl*,*k*_, *c*_*t*,*k*_, *d*_*f*,*k*_ and *e*_*c*,*k*_ correspond to the loading vectors for component *k,* for each source layer, target layer, time point, frequency point, and between-area connection. Loading vectors were estimated using alternating least square with random initialization and a non-negativity constraint. The variance of the data is usually kept in one of the loading vectors of PARFAC model and the other loading vectors are normalized to have variance of 1. In our case, the variance was kept in the temporal loadings. We used the N-way Matlab toolbox for PARAFAC decomposition ^62^.

To derive the most informative and valid PARAFAC model, an appropriate number of components needs to be selected. With too few components, the true underlying components of the data cannot be extracted, while with too many, the results will contain correlated components that do not represent the underlying variables. To identify the appropriate number of components, we combined multiple indicators of model quality and visual inspection of the resulting components, following previous works ^26–30^. To exclude the possibility that component selection was driven by data from a subset of animals, we first created bootstrapped averages by randomly selecting 8 (out of 11) animals. For each bootstrapped average (n=10), we calculated Corcondia scores, variance explained, mean squared error, and model convergence. Using these scores across bootstraps we found that models with 4 components were the best choice for our data: they substantially increased model fit as compared to sparser models, while including more components resulted in high correlations between some of them.

Finally, to verify the reliability of the PARAFAC model across animals, we bootstrapped iPDC averages (n=500, random selection as above) and extracted the four components for each bootstrap. We checked component consistency across bootstraps by calculating the pairwise correlations between the loading vectors of PARAFAC components, resulting in intra- and inter-component consistency values.

### Statistical analysis on PARAFAC loadings

To specify the characteristic of each network, we investigated the loading vectors of the PARAFAC models using statistical analysis. To indicate the source and target layers that contribute most to each network, we compared the loading values of each source/target layer over bootstraps to uniform distribution of weights over layers using t-test. Uniform distribution indicates no laminar specificity for the network, while layers with loading values larger than the uniform distribution have significant contribution to that network.

For temporal loadings, we determined significant evoked responses from baseline by comparing the temporal loading values of each post-stimulus time point with averaged pre-stimulus temporal loading values, using t-test and Bonferroni correction. For visualization purpose, we converted the temporal loadings to percentage of change by subtracting and then dividing each temporal loading by its averaged pre-stimulus loading values.

Since the variance of the data is preserved in the temporal loadings, we indicated the relative overall amplitude of each PARAFAC component by dividing its averaged temporal loading by the sum of averaged temporal loadings of all components.

### Model comparison of frequency distributions

To characterize frequency distribution of the PARAFAC components, we fitted three distributions to the frequency loadings in each bootstrap separately. Power-law (in the form of *c*. *f*^−*β*^), lognormal (log *N*(*μ*, *σ*)), and exponential (*c*. *e*^**β**.*f*^) distributions were fitted using *nlinfit* function in Matlab and the mean squared error for each fitting was calculated. The model with the lowest average mean squared error over the bootstraps was considered as the best model fit.

We also fitted the same models to the power spectrum density of LFPs, calculated from the AR coefficients, for each bootstrap and each ROI separately and used mean squared error to indicate the best model fit.

### Functional hierarchy analysis

We used loading vectors of between-area connectivity strengths to calculate hierarchy scores, employing a method specifically proposed for directed functional connectivity strengths ^12^. First, the directed influence asymmetry index (DAI_ij_) was estimated based on between-area loading vector with element *e*_*ij*_ indicating the connection from area *j* to *i*.

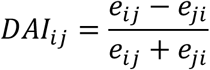

We scaled DAIs to the range [-2.5 2.5] in order to allow 6 levels of hierarchy (considering 6 areas). Then, for each target area, we shifted the DAIs from all source areas so that the smallest value was 1. Then we averaged the rescaled and shifted DAIs 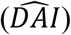 for each source to estimate the hierarchy score of the source area as 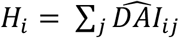.

### Receptive field analysis

The significant receptive fields (RF) were calculated based on a permutation-based method used in ^2^ and using the strength of evoked LFPs in a Gabor stimuli condition in the dataset. RF distances were calculated as the Euclidean distance between the centroids of the significant RFs. We only considered the between-area connections which had source and target areas with significant receptive fields (80 %).

To examine the relationship between RF distance and functional connectivity strengths, we reconstructed between-area connectivity strengths from the PARAFAC results using Kronecker products of the loading vectors, for each component and bootstrap separately. We then averaged the reconstructed matrix over time, frequency and layers to obtain a single connectivity value for each between-area connection.

Using linear mixed-effect regression, we modelled connectivity strengths as a function of the continuous predictor RF distance, including bootstrap and between-area connection as random effects over the intercept, to account for between-area and animal variability respectively. This model was preferred over a simpler fixed-effects model without random effects, as shown by a Likelihood Ratio Test. Modeling was done using the *fitlme* function with maximum likelihood estimation in Matlab.

## Acknowledgement

This research was supported by the Swiss National Science Foundation (grants PP00P1_157420 and PP00P1_183714).

## Author contributions

EB and GP: Investigation, Methodology, Writing; EB: Data curation, Formal Analysis, Visualization; GP: Conceptualization, Funding acquisition, Supervision.

## Competing Interests statement

The authors declare no competing financial or non-financial interests.

## Notes

### Competing Interest Statement

The authors have declared no competing interest.

